# Transcriptional architecture underlying development, adaptation, and domestication in Northern Wild Rice (*Zizania palustris* L.)

**DOI:** 10.64898/2026.03.30.715309

**Authors:** Maybell DM. Banting, Matthew Haas, Sapphire Coronejo, Lillian McGilp, Laura M. Shannon, Jennifer Kimball

## Abstract

*Zizania palustris* (Northern Wild Rice) is an aquatic grass native to North America and a crop wild relative of *Oryza sativa* with ecological, cultural, and agricultural significance. Here, we present a transcriptomic atlas spanning 20 tissues across six major developmental stages. Seed tissues showed shifts in abscisic acid, gibberellin, and ethylene pathways that define the hormonal basis of deep dormancy in this recalcitrant species and its release. Leaf development showed stage-specific reprogramming, from hypoxia-responsive programs in submerged tissues to cell wall remodeling and redox regulation during aquatic–aerial transitions, with photosynthesis and carbohydrate metabolism defining flag leaves and *SRG1* divergence emerging as a defining feature of leaf development in *Z. palustris*. Reproductive tissues expressed duplicated homologs of well-recognized shattering genes with divergent regulation, consistent with subfunctionalization after whole-genome duplication. These findings provide new insight into traits underlying ecological adaptation and domestication in *Z. palustris* and related grasses.

**CORE IDEAS:**

- A first transcriptomic atlas for Northern Wild Rice maps gene expression across 20 tissues and six developmental stages, providing a foundational functional genomics resource for *Zizania palustris*.
- Seed dormancy and release are driven by coordinated hormone reprogramming, with shifts in ABA, GA, and ethylene pathways across whole seed, embryo, and endosperm.
- Leaf development is defined by aquatic-to-aerial transcriptional transitions, moving from hypoxia-responsive programs in submerged tissues to cell wall remodeling, redox regulation, and photosynthesis/metabolism in aerial and flag leaves.
- Whole-genome duplication underlies regulatory divergence in domestication traits, as duplicated homologs of canonical seed shattering genes show paralog- and tissue-specific expression consistent with subfunctionalization.
- The atlas identifies tissue-specific modules and stable housekeeping candidates, enabling improved experimental design (e.g., expression normalization) and accelerating candidate prioritization for breeding, GWAS/eQTL, and trait discovery.

**PLAIN LANGUAGE SUMMARY:** Northern Wild Rice (*Zizania palustris*) is an aquatic grass native to North America that is important for ecosystems, agriculture, and Indigenous cultures. In this study, we created a detailed gene expression map across 20 different tissues and six stages of plant development. We found that seeds strongly regulate plant hormones that control dormancy, helping explain why Northern Wild Rice seeds remain dormant for long periods and how they eventually germinate. Leaves showed clear changes in gene activity as plants moved from underwater growth to above-water growth, shifting from low-oxygen stress responses to processes that support photosynthesis and structural strength. In flowers, we identified duplicated genes involved in seed shattering that are regulated differently, likely due to past genome duplication events. Together, these results improve our understanding of how Northern Wild Rice is adapted to aquatic environments and how key traits relevant to domestication have evolved.

## 1 Introduction

Understanding how plants adapt and thrive in rapidly changing environments is a central challenge in plant biology, with direct implications for food security, ecosystem stability, and conservation. *Zizania palustris* (Northern Wild Rice; NWR), a culturally and ecologically significant grain-producing aquatic grass native to North America’s wetlands, offers a unique model for dissecting the genetic and molecular basis of adaptation to aquatic habitats and seasonal variability. As a crop wild relative (CWR) of rice (*Oryza sativa*) within the Oryzeae tribe, *Z. palustris* has followed a distinctive evolutionary trajectory, diverging from the Oryza lineage approximately 20–30 million years ago (Y. Guo and Ge, 2005; Haas et al., 2021), and has since evolved a suite of adaptive traits that enable persistence in cold-adapted dynamic freshwater ecosystems.

Despite its importance as both a crop and a species providing important ecosystem services, *Z. palustris* largely remains genetically undercharacterized. Recent advances in genomics have produced high-quality reference genomes and identified ancient whole genome duplication (WGD) events in the *Zizania* lineage (L. Guo et al., 2015; Haas et al., 2021; Yan et al., 2022), but functional genomic resources have lagged behind. This gap limits our understanding of the regulatory networks underlying developmental transitions, stress resilience, and traits critical for the continued domestication of the species, such as seed dispersal and dormancy, which are especially relevant as wetland ecosystems face accelerating climate change.

Here, we present the first comprehensive transcriptomic atlas of *Z. palustris*, spanning 20 tissue types and all major developmental stages of its annual life cycle. This study focuses on the regulation of seed dormancy release, leaf developmental transitions, reproductive differentiation, and seed shattering, traits that define the life cycle and domestication of *Z. palustris*. By integrating high-resolution gene expression profiling with comparative genomics, we (i) infer transcriptional transitions associated with key biological processes, (ii) identify expression signatures linked to domestication traits such as seed shattering, and (iii) define tissue-specific and broadly expressed gene modules. Our dataset advances functional genomics in a foundational grain-producing aquatic grass, and provides a critical resource for trait discovery, crop improvement, and conservation genomics within the Oryzeae tribe.

## 2 Materials and Methods

### 2.1 Plant Materials

We used the cNWR variety ‘Itasca-C12’, previously employed in the construction of the *Zizania palustris* reference genome (NCBI database, GCA_019279435.1) (Haas et al., 2021). This genotype, maintained by the University of Minnesota’s (UMN) cNWR breeding program, exhibits intermediate seed recalcitrance that precludes conventional seed bank preservation (Pence, 1995). The individual plant (∼15% heterozygosity) from which the reference genome was derived has been propagated in an open-pollinated environment by growing ∼300 seeds per cycle to retain heterozygosity. Post-harvest seeds were stored in distilled water at 1–3°C in the dark for approximately one year (McGilp et al., 2023).

In winter 2022, 160 plants were grown in controlled conditions at the UMN Plant Growth Facilities (St. Paul, MN). We collected twenty distinct tissue types spanning leaves, stems, roots, male and female floral organs, and seeds across major developmental stages. Each tissue was sampled with two to three biological replicates (Table S1). Developmental staging followed the principal phenological stages (Duquette and Kimball, 2020) and included seed maturation, dormancy, germination, early and late booting, leaf emergence stages (submerged, floating, aerial, flag), and anthesis. Each biological replicate represents an independent plant sampled at the same developmental stage and tissue type, with all replicates processed independently through RNA extraction, library preparation, and sequencing.

### 2.2 RNA Extraction and Sequencing

RNA was extracted from each tissue sample using a Qiagen RNeasy kit (Qiagen, Germantown, Maryland). Following extraction, the quality and quantity of the RNA was assessed using a Nanodrop spectrophotometer (Thermo Fisher Scientific, Wilmington, Delaware). The extracted RNA samples were stored at -80°C to preserve their stability. After storage, the samples were sent to the University of Minnesota Genomics Center (UMGC, http://genomics.umn.edu) for library preparation and sequencing, using a 56 unique dual-indexed (UDI) approach with the Illumina TruSeq stranded mRNA kit. Sequencing was performed using 150-bp paired-end reads on an Illumina NovaSeq platform.

### 2.3 Gene Atlas and Reference-Guided Transcriptome Assembly

Gene expression estimates were produced and summarized using a modified pipeline based on methodologies from recent related transcriptome studies (Machado et al., 2020). The RNA-seq data were analyzed using the Tuxedo protocol (Pertea et al., 2015), incorporating HISAT2 v.2.2.11 for alignment (Kim et al., 2015) and StringTie v.2.2.1 for transcript assembly (Pertea et al., 2015). Initial quality assessment of the raw sequence data was conducted using FastQC v.0.11.9 (Andrews, 2010). Subsequently, low-quality reads and adapter contamination were removed using Trimmomatic v.0.39 (Bolger et al., 2014). The strandedness of the RNA-seq data was determined using the how_are_we_stranded_here tool v.1.0.1 (Signal & Kahlke, 2022). To eliminate rRNA reads, the BBDuk tool from the BBTools suite (Bushnell, 2014) was utilized, employing a kmer length of 27 as a filtering threshold and a compressed version of the SILVA database v.38.1 for decontamination (Quast et al., 2012). The quality of the trimmed and filtered reads was reassessed using FastQC v.0.11.9. These high-quality reads were then aligned to the *Z. palustris* reference genome (Itasca-C12; GCA_019279435.1; ASM1927943v1) (Haas et al., 2021), which was obtained from NCBI GenBank (NCBI, https://www.ncbi.nlm.nih.gov). Alignment was performed using HISAT2, followed by sorting of the aligned reads by chromosome position, conversion to BAM format, and indexing using SAMtools v.1.16.1 (Danecek et al., 2021). The sorted BAM files served as the basis for transcript assembly and abundance estimation, performed using StringTie (Pertea et al., 2015), with transcript abundances quantified in transcript per million mapped reads (TPM). These abundances were then translated into gene-level read counts using the prepDE.py script included with StringTie. Unless stated otherwise, all programs were used with their default parameters.

### 2.4 Differential Gene Expression Analysis

Gene-level read counts were imported into R using the tximport package (Soneson et al., 2015) and normalized using a variance-stabilizing transformation. Differential expression analysis was performed with DESeq2 (Love et al., 2014). P-values were adjusted for multiple testing using the Benjamini-Hochberg false discovery rate procedure as implemented in DESeq2 and reported as adjusted p-values (padj). By default, DESeq2 applies independent filtering based on mean normalized counts and flags extreme count outliers using Cook’s distance, resulting in adjusted p-values of NA for filtered genes (Love et al., 2014). Genes with nonzero counts across all samples were retained for analysis. Samples were grouped into “untreated” (control) and “treated” categories to enable pairwise comparisons across tissues and developmental stages. Differentially expressed genes (DEGs) were defined by an absolute log₂ fold change ≥ 2 and an adjusted p-value < 0.05.

To assess global expression patterns, evaluate concordance among biological replicates, and identify potential confounding effects, we performed multivariate analyses including Principal Component Analysis (PCA) and hierarchical clustering on variance-stabilized expression values. PCA was performed using *plotPCA()* function from DESeq2 (Love et al., 2014) and visualized the results using *ggplot2* v3.5.1 (Wickham, 2016). Additionally, pairwise Pearson correlation coefficients were calculated from normalized expression values to quantify transcriptional similarity across tissues. These coefficients were transformed into a distance matrix and subjected to hierarchical clustering using the *hclust()* function in R. The resulting dendrogram and heatmap, visualized with the pheatmap package (Kolde, 2019), revealed clustering patterns that reflected tissue type, developmental stage, and inferred biological function.

### 2.5 Functional Annotation and Gene Ontology Enrichment

To investigate the functional landscape of the *Zizania palustris* transcriptome, we performed a multi-tiered analysis comprising transcriptome-wide functional annotation and Gene Ontology (GO) enrichment. Coding sequences were first predicted using TransDecoder v5.7.1 (Haas, 2019), enabling the delineation of protein-coding regions from non-coding transcripts in the assembled transcriptome. The resulting peptide sequences were annotated through homology-based searches using HMMER v3.4 (Eddy, 2011) against the Pfam database (release 37.1) (Mistry et al., 2021) and BLASTP searches against the NCBI non-redundant (NR) protein database (E-value threshold of 1×10−10). In addition, InterProScan (Jones et al., 2014) was used to assign protein domain-based annotations and consolidate results from multiple functional databases, including Pfam, PANTHER, and SMART (Letunic et al., 2021; Mi et al., 2021; Mistry et al., 2021).

GO enrichment analysis was conducted using the topGO R package (Alexa and Rahnenfuhrer, 2016). Enrichment was tested using Fisher’s exact test with the classic algorithm, with Biological Process (BP), Molecular Function (MF), and Cellular Component (CC) categories analyzed separately. The classic algorithm evaluates GO term enrichment without applying global multiple-testing correction, reflecting the hierarchical dependency structure of the GO. Raw p-values were used to identify enriched GO terms, and terms with p-values < 0.01 were retained for downstream interpretation (Alexa and Rahnenfuhrer, 2016).

### 2.6 K-means Clustering of Gene Expression Profiles

Gene co-expression modules were identified by k-means clustering of variance-stabilized RNA-seq expression values. Transcript-level TPMs were filtered to retain genes with TPM ≥ 1 in at least one tissue and collapsed to gene-level by summing across isoforms. Variance stabilization was performed using DESeq2 (Love et al., 2014). The optimal number of clusters was determined using a gap statistic with 500 bootstraps and 25 random starts, implemented in factoextra v1.0.7 (Kassambra and Mundt, 2020). This analysis supported k=19, which was used for k-means clustering with a fixed random seed. Expression values were averaged across replicates and visualized using ggplot2 v3.5.1 (Wickham, 2016).

### 2.7 Identification of Tissue Specific and Broadly Expressed Genes

To quantify tissue-specific gene expression, we calculated the Tau index (τ) using log2-transformed TPM values averaged across biological replicates for each tissue. The τ index ranges from 0 (broad expression) to 1 (tissue-specific expression). The τ index for each gene was computed using the following formula (Machado et al., 2020):

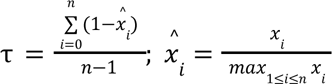

where, *x_i_* represents the expression level of the gene in the *i* tissue, and *n* denotes the total number of tissues analyzed.

Following the classification criteria as previously described (Lüleci and Yılmaz, 2022), genes were grouped into four expression categories based on their expression levels and τ indices. Genes with TPM < 1 across all tissues were classified as exhibiting null expression, indicating no detectable activity. Genes with TPM < 5 in all tissues were considered weak expression, reflecting low overall expression levels. Broadly expressed genes were defined as those with TPM > 5 and τ < 0.85, indicating widespread expression across multiple tissues. Tissue-specific genes, on the other hand, were characterized by TPM > 5 and τ ≥ 0.85, indicating a strong preferential expression in one or a limited subset of tissues.

Housekeeping (HK) genes, characterized by consistent expression across diverse conditions, were identified using previously established criteria (Machado et al., 2020). To be identified as an HK gene, a gene had to be expressed in all samples, ensuring its presence across tissues. The mean expression level for each gene was calculated as the average TPM value across all samples. To evaluate expression variability, the coefficient of variation (CoV) was computed by dividing the standard deviation of expression levels by the mean. Additionally, the ratio of maximum to minimum fold change (MFC) was calculated by dividing the highest TPM value by the lowest. These two metrics were combined into a product score (MFC-CoV) by multiplying the CoV and MFC values for each gene. Genes with MFC-CoV scores falling within the first quartile, representing the least variable and most stable expression profiles, were classified as HK genes. To further validate their broad expression and lack of tissue specificity, these genes were analyzed using τ index.

## 3 Results

### 3.1 Quantitative Overview of the Northern Wild Rice Transcriptome

To establish a comprehensive transcriptomic atlas for *Z. palustris*, we generated RNA-seq data from 55 samples representing 20 tissues spanning major developmental stages, including seed, leaf, stem, root, and panicle tissues (Figure S1; Table S1). Sequencing generated ∼1.8 billion high-quality paired-end reads, with an average alignment rate of 96.23% to the *Z. palustris* reference genome, equating to ∼33.8 million aligned reads per sample (Data S1). Using criterion of TPM ≥ 1 in at least one tissue, 38,777 genes were identified as expressed (Table S1), compared to 46,491 putative protein-coding genes in the first *Z. palustris* transcriptome assembly (Haas et al., 2021). Genes were further categorized by expression levels, with low-to-moderate expression identified most commonly in ripe and dormant seeds and female panicles (TPM ≥ 1 & < 10), whereas stems and panicles had the greatest proportion of highly expressed genes (TPM ≥ 10).

Heirarchical clustering and principal component analysis (PCA) both identified two major clades separating seed-related tissues from vegetative and reproductive organs. (Figure 1A, B). In the PCA, this separation explained 78.43% of the variance along PC1, while PC2 (6.12% variance) distinguished variation across developmental stages within the two clades. Within the seed clade, samples clustered along a developmental gradient beginning with ripe seed (beginning dormancy), then shifting to dormant seed, embryo, and endosperm samples, followed by stratified (non-dormant) seed, embryo, and endosperm samples, and finally germinating seeds (Figure 1B). With the second major clade, clustering began with roots, followed by coleoptile, submerged leaves, then transitioned to a mixture of early-, mid-, and late-boot stem tissues, followed by a male and female panicle cluster and finally, the aerial leaves. Pairwise differential expression between tissues identified 26,115 unique differentially expressed genes (DEGs) (log₂FC ≤ –2 or ≥ 2; adjusted p < 0.05), with the highest number of DEGs between seed and other tissue types (Figure 1C).

**Figure 1.**
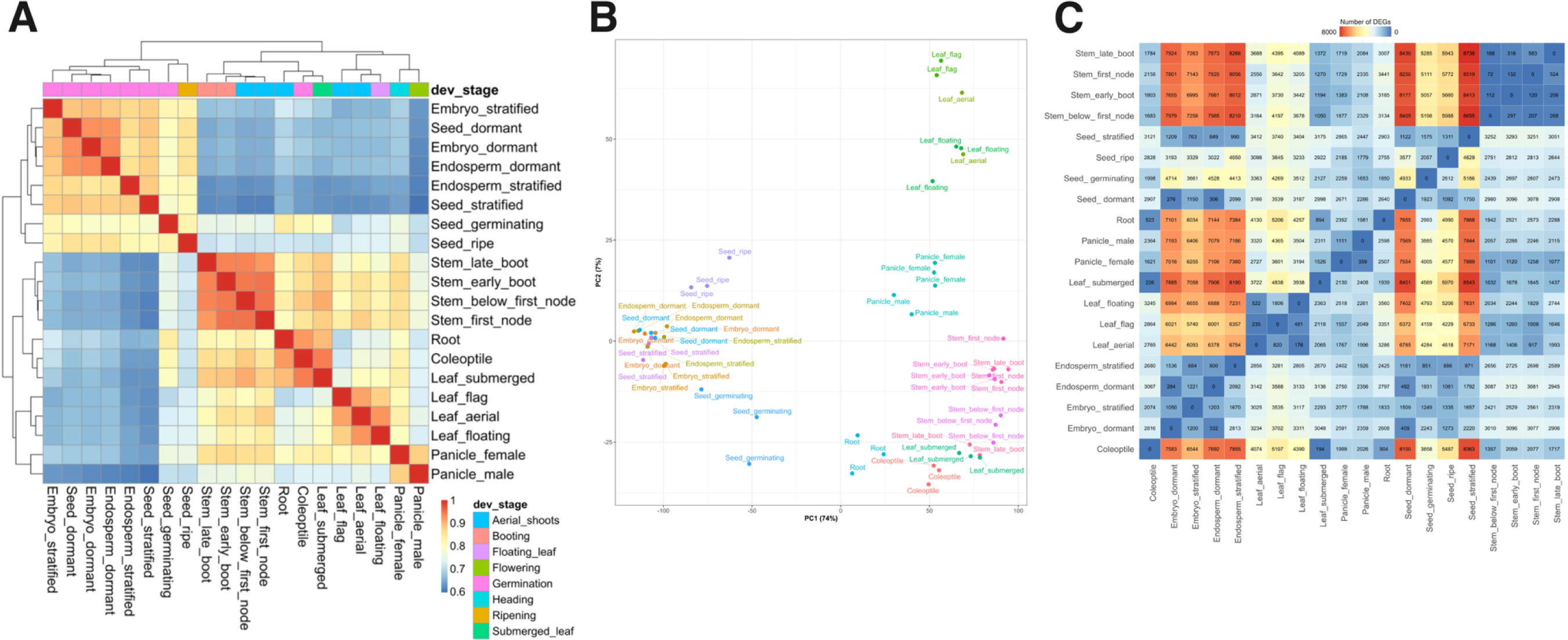
Global transcriptome relationships and differential expression across *Z. palustris* tissues. (A) Heatmap of pairwise Pearson correlation coefficients (R) among 20 tissue samples based on variance-stabilized gene expression values. Hierarchical clustering reveals strong tissue-specific transcriptomic signatures, with closely related tissues clustering together. (B) Principal component analysis (PCA) of transcriptomes from 20 distinct tissues. Each point represents a biological sample, color-coded by tissue type. PC1 explains 78.43% of the total variance and separates major tissue groups, while PC2 accounts for 6.12% of the variance and captures finer-scale expression differences. (C) Matrix of tissue-by-tissue differential expression comparisons. The upper triangle displays the number of genes upregulated in the row tissue relative to the column tissue, while the lower triangle shows the number of downregulated genes in the same comparison.

### 3.2 K-means Clustering Identifies Tissue-Specific and Housekeeping Gene Modules

To further resolve transcriptional organization, we applied k-means clustering to variance-stabilized expression data, which grouped expressed genes into 19 clusters reflecting distinct tissue- and stage-specific programs (Figure 2A; Data S2), with the number of clusters supported by gap statistic analysis. Seed-associated clusters (10–12) captured transcriptional states corresponding to sequential developmental transitions from seed filling to dormancy (Data S2). Cluster 12 was significantly enriched for GO terms related to seed, fruit, and reproductive structure development; cluster 10 for nutrient reservoir, glycogen synthase, and glucosyltransferase activity; and cluster 11, with higher expression in dormant embryo and endosperm tissues, for chromatin compaction processes. Vegetative-associated clusters (13, 14, 16, 17, and 19) displayed moderate to high expression in leaf and stem tissues with enrichment for oxidative stress responses as well as photosynthetic and light-response functions. Cluster 17 was strongly expressed in submerged leaves, coleoptiles, and early-stage stem tissues with enrichment for translation, ribosome biogenesis, and rRNA processing. Panicle-specific programs were represented by cluster 4, enriched for isoprenoid and steroid biosynthesis. Cluster 18 displayed uniformly high expression across tissues, enriched for ribosomal and translational functions, consistent with housekeeping activity.

**Figure 2.**
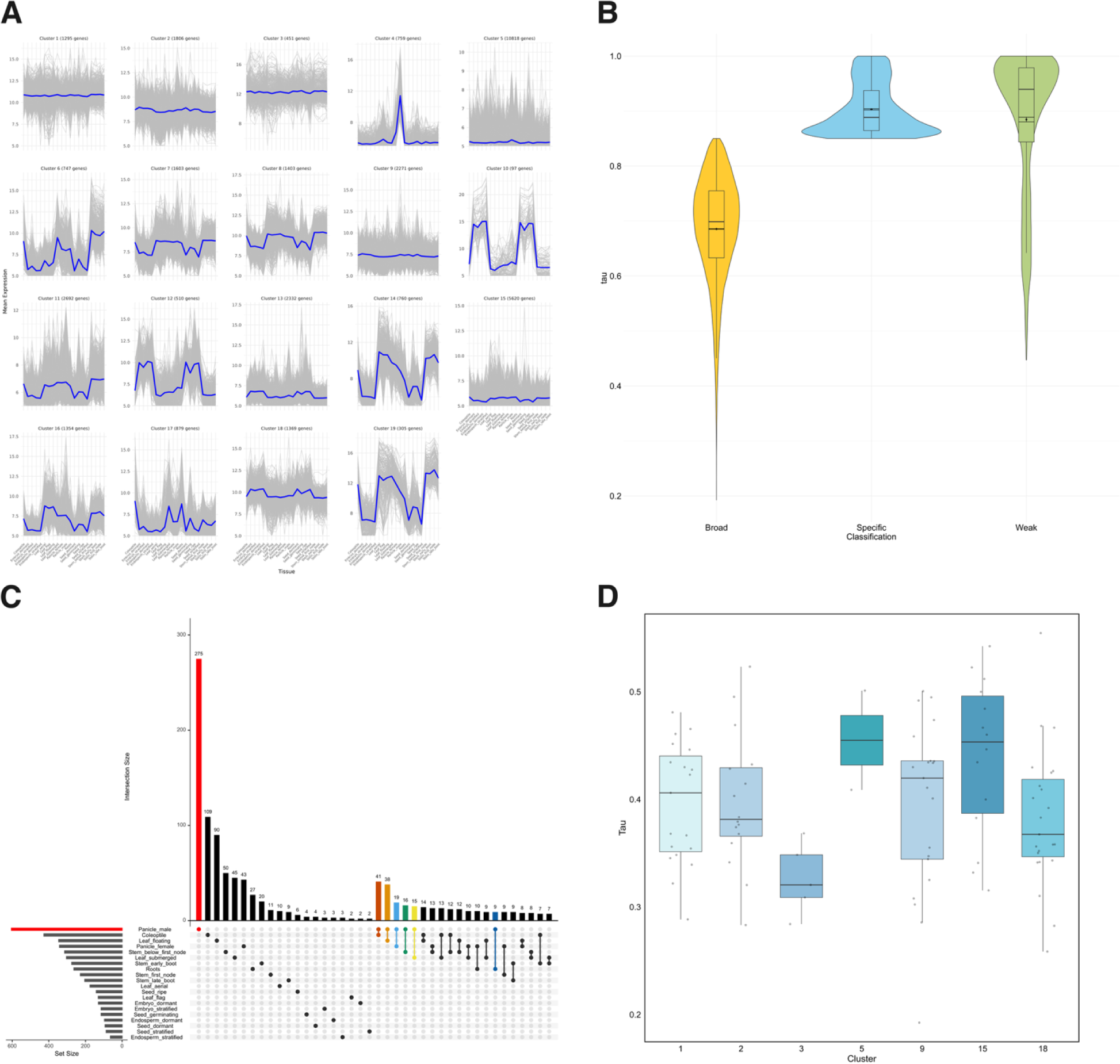
K-means clustering, tissue-specific genes, and housekeeping candidates in *Z. palustris*. (A) K-means clustering of all expressed genes in 19 clusters based on variance-stabilized expression values across 20 tissues. Each panel represents one cluster, with individual gene expression profiles in grey and the cluster centroid in blue. (B) Distribution of the tau index (τ) for genes classified as broad, specific, or weak. (C) UpSet plot illustrating intersection patterns among tissue-specific genes. Bar heights represent the number of genes specific to individual tissues (bottom left) or their combinations (top). Highlighted intersections include genes specific to Panicle_male alone (red) and in combination with other tissues (colored bars). (D) Boxplot of τ values for putative housekeeping genes assigned to k-means clusters with low expression variability (clusters 1, 2, 3, 5, 9, 15, and 18).

To independently validate tissue-specificity, we calculated the tau (τ) index for all 38,777 expressed genes (Figure 2B and Data S3). Approximately 1,926 genes (5%) were highly tissue-specific (τ ≥ 0.85), with the largest numbers in male panicles (604), coleoptiles (427), and floating leaves (346), whereas dormant and stratified seeds had the fewest (Figure 2C and Figure S2). Conversely, 767 genes were expressed in all tissues (TPM ≥ 1) and identified as candidate housekeeping genes. Ranking these by expression stability using the coefficient of variation × maximum fold change (CoV × MFC) identified 192 robust candidates, most concentrated in cluster 18 (Figure 2D and Data S4).

Together, these analyses define the transcriptional landscape of *Z. palustris*, revealing clear distinctions between seed, vegetative, and reproductive tissues, stage-specific programs within seed development, and tissue-enriched modules in both submerged and aerial organs. Integration of clustering, tau index, and stability metrics provided a robust framework for identifying both tissue-specific regulatory modules and broadly conserved gene sets for functional studies and expression normalization.

### 3.3 Developmental and Hormonal Reprogramming During Seed Dormancy Transitions

Pairwise comparisons among developmental stages in *Z. palustris* identified different numbers of DEGs. In whole seeds, 2,865 DEGs distinguished dormant from stratified stages, while 6,727 DEGs separated stratified from germinating seeds, and 4,647 DEGs marked the transition from ripening to dormancy (Figure 3A). These comparisons revealed progressive layers of regulation: dormancy onset was associated with chromatin remodeling, stratification with modest but targeted metabolic adjustments, and germination with broad metabolic activation.

**Figure 3.**
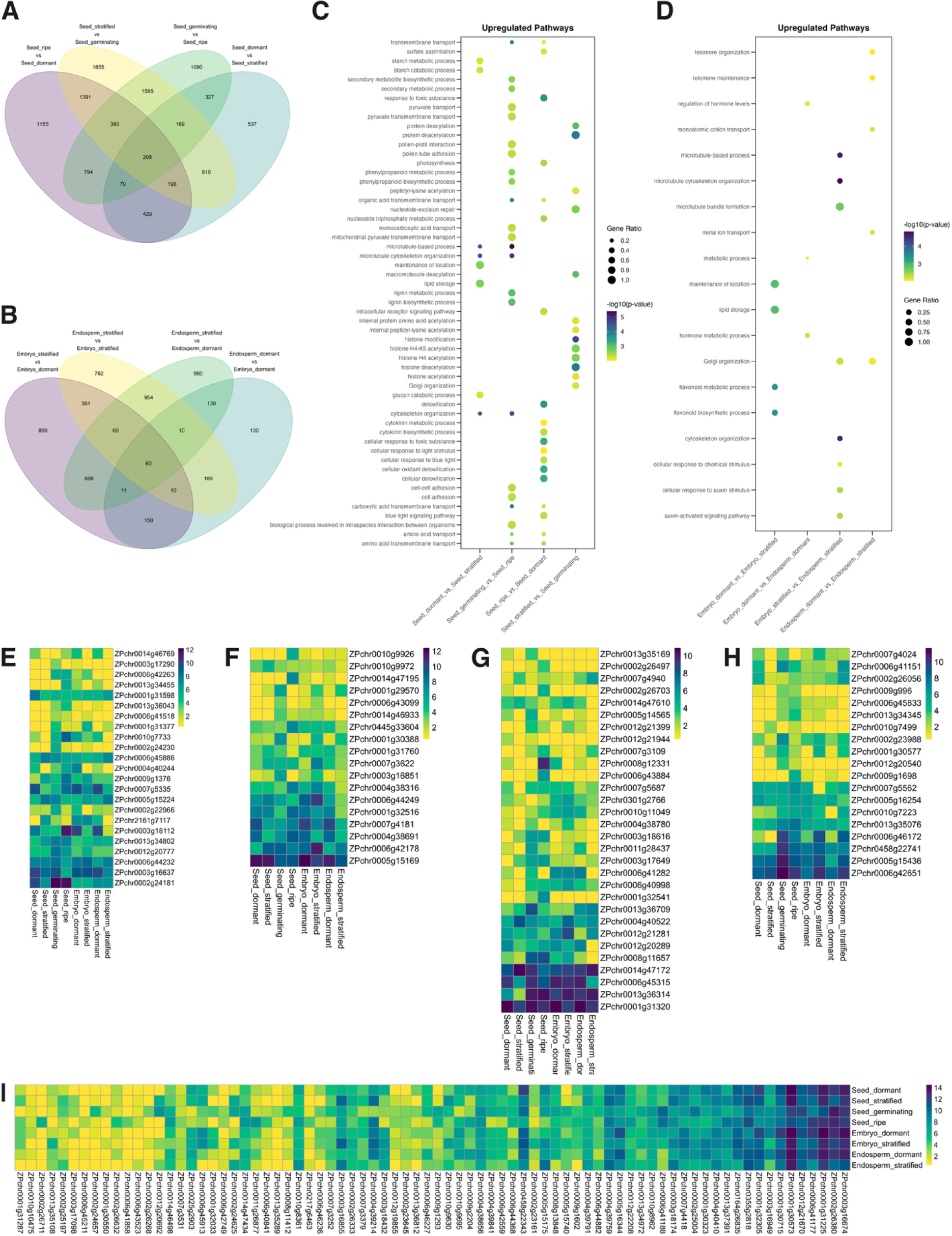
Differential gene expression, functional enrichment, and hormone-associated transcriptional patterns during seed dormancy transitions in *Z. palustris*. (A) Venn diagram showing overlaps among differentially expressed genes (DEGs) in whole-seed contrasts: ripe vs. dormant, stratified vs. germinating, germinating vs. ripe, and dormant vs. stratified. (B) DEG overlaps in embryo and endosperm tissue contrasts: dormant embryo vs. stratified embryo vs. dormant endosperm, stratified embryo vs. stratified endosperm, and dormant endosperm vs. stratified endosperm. (C) Gene Ontology enrichment among upregulated DEGs in whole seeds, (D) embryo and endosperm tissues. (E–I) Heatmaps of DEGs related to (E) abscisic acid, (F) gibberellin, (G) cytokinin, (H) jasmonic acid, and (I) ethylene pathways. Expression values are variance-stabilized counts (DESeq2). Genes shown are differentially expressed in at least one contrast across the whole seed, embryo, or endosperm.

Gene ontology (GO) enrichment analysis of upregulated DEGs revealed distinct biological processes underlying each developmental transition (Figure 3C). In ripe versus dormant seeds, 29 enriched GO terms were associated with nucleosome assembly, histone modification, and DNA packaging, indicating chromatin-level changes during dormancy onset. The transition from dormancy to stratification exhibited limited enrichment (6 GO terms), including carbohydrate metabolism, cell wall reorganization, and oxidoreductase activity, consistent with limited but targeted metabolic changes during cold stratification. In contrast, the shift from stratified to germinating seeds was marked by the most extensive changes, with 37 enriched GO terms related to translational activation, ribosome assembly, protein folding, RNA metabolism, and mitochondrial gene expression. These processes reflect a broad reactivation of cellular metabolism required for growth. The germinating versus ripe comparison revealed eight enriched GO terms related to photosynthesis, secondary metabolism, and sulfur compound metabolism. Several categories, including RNA processing, protein refolding, and stress response, were enriched across multiple contrasts, suggesting shared transcriptional signatures of seed transition states.

Embryo and endosperm tissues followed distinct developmental trajectories. At the dormant stage, embryos and endosperms differed by 610 DEGs with a single enriched term, but stratification expanded these differences to 2,346 DEGs and eight enriched terms, demonstrating strong tissue-specific reprogramming (Figure 3B,D). Within embryos, 2,250 DEGs separated stratified from dormant stages, enriched for flavonoid biosynthesis, lipid storage, telomere organization, and hormone regulation, consistent with activation of developmental and protective pathways (Figure 3B,D). Endosperms underwent even broader changes, with 2,883 DEGs distinguishing stratified from dormant tissues, enriched for mitochondrial protein import, translation initiation, and chromatin organization, reflecting large-scale metabolic and regulatory restructuring (Figure 3B,D). These results indicate that embryos and endosperms contribute jointly to dormancy release but through complementary transcriptional programs, with embryos emphasizing developmental control and endosperms emphasizing metabolic reprogramming.

To elucidate hormone-mediated regulation during dormancy transitions, we analyzed DEGs annotated to biosynthesis, signaling, and perception pathways of five major phytohormones: abscisic acid (ABA), gibberellin (GA), cytokinin (CK), jasmonic acid (JA), and ethylene (Figure 3E–I; Table S2-6). In total, 34 ABA-related, 29 GA-related, 22 CK-related, 25 JA-related, and 81 ethylene-related DEGs were identified across seed tissues and developmental stages. The expression dynamics of these hormone-associated genes aligned closely with physiological transitions.

ABA-related genes, including biosynthetic enzymes (*NCED*), perception proteins (PYR/PYL receptors), and signaling intermediates (PP2Cs, ABFs), were highly expressed in dormant tissues but progressively downregulated during stratification and germination. This decline indicates reduced ABA signaling during dormancy release (Figure 3E; Table S2). In contrast, GA-related genes, including biosynthetic (*GA20ox, GA2ox*), perception (GID1 receptors), and signaling (DELLA repressors) were upregulated during stratification and germination (Figure 3F; Table S3), consistent with GA’s established role in promoting germination.

CK-related DEGs, including biosynthetic (*IPT*) and degradation (*CKX*) enzymes along with type-B ARR transcription factors, showed their highest expression in stratified embryos, suggesting a role in embryo-specific regulation (Figure 3G; Table S4). JA-related genes, including biosynthetic enzymes (*AOC, OPR3*) and signaling regulators (COI1, JAZ proteins), were most highly expressed in dormant and stratified embryos (Figure 3H; Table S5), suggesting potential roles in dormancy maintenance or stress response. Ethylene-related DEGs, including biosynthetic (*ACS8, ACO1*), perception (ETR1), and downstream signaling components (ERF1, EIN3), were upregulated in germinating tissues (Figure 3I; Table S6), reflecting ethylene’s contribution to seedling emergence and growth.

These patterns demonstrate that seed dormancy release in *Z. palustris* involves coordinated developmental reprogramming and dynamic regulation of hormone biosynthesis and signaling. ABA signaling is progressively repressed, GA and ethylene pathways serve as central regulatory shifts, CK and JA contribute tissue-specific functions, particularly in the embryo.

### 3.4 Stage-Resolved Transcriptomic Shifts Define the Developmental Progression of Leaves

Leaf development in *Z. palustris* proceeds through a series of morphologically and physiologically distinct stages, beginning with submerged juvenile leaves and culminating in fully mature flag leaves at the reproductive stage. To characterize the transcriptional programs underlying this development, we profiled gene expression across four distinct leaf tissue types, including submerged, floating, first aerial, and flag leaves. These tissues represent sequential developmental transitions that differ both in environmental context (aerobic versus anaerobic growth) and in maturity (juvenile versus reproductive). In presenting these results, we highlight representative genes whose functions in hormone signaling, cell wall remodeling, or metabolism illustrate the major regulatory and structural changes occurring at each developmental stage.

The most extensive reprogramming occurred between submerged and floating leaves, with 6,288 DEGs including 3,931 more highly expressed in submerged tissues (Figure 1C, 4A). The floating-to-aerial transition involved 694 DEGs and the aerial-to-flag transition 1,055 DEGs (Figure 1C, 4B,C). Only 79 DEGs were shared across all three contrasts (Figure 4D), showing that each stage is marked by a distinct expression profile. These contrasts define a developmental trajectory in which the major transcriptional reprogramming occurs early, during the submerged-to-floating transition, followed by more targeted remodeling as leaves adapt to aerial growth and finally a shift toward structural and metabolic maturation in the flag leaf.

**Figure 4.**
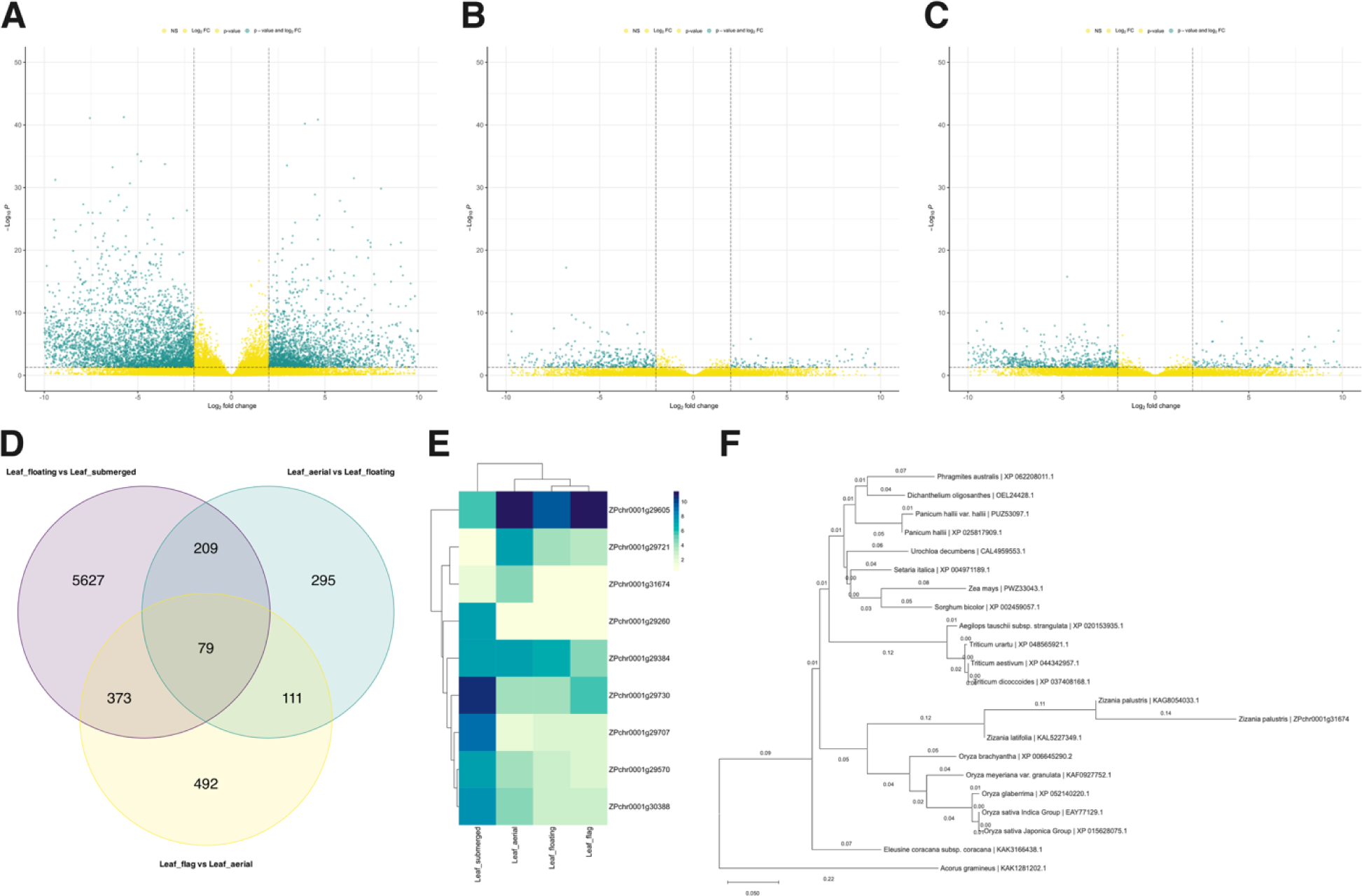
Transcriptome changes during leaf development and evolutionary analysis of an *SRG1* homolog in *Z. palustris* (A-C) Differential expression analysis between developmental leaf stages using DESeq2. Volcano plots show pairwise comparisons for Leaf_floating vs Leaf_submerged (A), Leaf_aerial vs Leaf_floating (B), and Leaf_flag vs Leaf_aerial (C). Genes meeting both thresholds (|log2 fold change| ≥ 2 and adjusted p-value < 0.05) are shown in teal; genes meeting only the fold change or p-value threshold are shown in yellow, respectively. (D) Venn diagram showing the number of differentially expressed genes (DEGs) that are shared and unique across three leaf contrasts. (E) Heatmap of variance-stabilized expression values for selected DEGs in the transition from submerged to floating leaves. (F) Phylogenetic tree constructed using the Neighbor-Joining method in MEGA12. The analysis included 22 protein sequences aligned using CLUSTALW. Evolutionary distances were calculated using the Poisson correction method with pairwise deletion of gaps, resulting in a final alignment of 451 positions. Bootstrap support values from 1,000 replicates are shown at internal nodes. The *Z. palustris* gene *ZPchr0001g31674* is included as an *SRG1* homolog.

The submerged-to-floating transition was enriched for 24 biological processes related to amino acid biosynthesis, chloroplast development, and light signaling (Table S7). Upregulated genes included *GIL1* (*ZPchr0001g29605*), a phytochrome-dependent regulator of gravitropism (Allen et al., 2006), and *RER3* (*ZPchr0001g29721*), which promotes mesophyll cell division during early leaf development (Pérez-Pérez et al., 2013) (Figure 4A,E). Hormone-related genes were repressed, including *GH3.1-like* (*ZPchr0001g29260*), which conjugates auxin to amino acids to reduce active IAA (Staswick et al., 2005), and *GA3ox2-like* (*ZPchr0001g29570, ZPchr0001g30388*), required for production of bioactive gibberellins (Mitchum et al., 2006). *EXTENSIN-like* (*ZPchr0001g29707*), GDSL esterase/lipase (*ZPchr0001g29730*), and alpha-amylase 3 (*ZPchr0001g29512*) were induced, indicating starch mobilization and cell wall modification during the aquatic-to-aerial shift (Figure 4a, e). Among wall-related genes, *EXPANSIN-A1* (*ZPchr0004g39708*) was strongly induced, consistent with wall loosening required for leaf elongation in rice (Cho and Kende, 1997).

The floating-to-aerial transition involved fewer DEGs and a single enriched GO term, tropism (Table S7). ABA perception declined through repression of *PYL4-like* (*ZPchr0001g31377, ZPchr0005g15224*), while cytokinin and auxin regulators including *CKX11-like* (*ZPchr0001g31476*), *ARF2* (*ZPchr0001g32049*), and *IAA18-like* (*ZPchr0005g14447*) were activated. Remodeling processes continued with induction of *TCP21-like* (*ZPchr0006g41156*), a homolog of Arabidopsis CHE that integrates circadian timing with transcriptional control (Pruneda-Paz et al., 2009). Developmental regulators were downregulated at this stage, including *RLT2* (*ZPchr0005g14930*), a chromatin-associated factor that maintains vegetative growth in Arabidopsis (G. Li et al., 2012), and *MYB4-like* (*ZPchr0001g29381*), a repressor of C4H in the phenylpropanoid pathway (Jin, 2000). Repression of *MYB4-like* likely enabled phenylpropanoid activity, supporting lignin deposition and flavonoid accumulation for structural reinforcement and photoprotection in aerial leaves. Whereas the floating-to-aerial transition relied on targeted remodeling of hormone and wall-associated pathways, the aerial-to-flag transition was characterized by broad suppression of structural growth programs and activation of metabolic genes.

The aerial-to-flag transition showed limited GO enrichment, with proteasome assembly the only enriched process (Table S7). Wall-associated genes were repressed, including *CESA4* (*ZPchr0001g30616)*, required for secondary wall cellulose biosynthesis in rice (Tanaka et al., 2003), *NAC73-like* (*ZPchr0001g30518*), homologous to Arabidopsis NAC-domain master regulators of secondary wall formation (Zhong et al., 2008), and *EXPANSIN-A4* (*ZPchr0001g31259*), marking a reduction in wall loosening activity that had been prominent at earlier stages (Cho and Kende, 1997). Metabolism-related genes were induced, including ATP-dependent phosphofructokinase (*ZPchr0001g32646*), which catalyzes a key regulatory step in glycolysis (Mustroph et al., 2007), and *GLOX1* (*ZPchr0001g29762*), a glyoxal oxidase linked to redox balance in Arabidopsis (Phan et al., 2011). These changes show that leaf maturation is accompanied by reduced investment in wall remodeling and increased activity in primary metabolism and redox regulation.

We identified *ZPchr0001g31674*, annotated as Senescence-Related Gene 1 (*SRG1*), as the most differentially expressed gene across all pairwise comparisons of leaf tissues. *ZPchr0001g29384* is also annotated as *SRG1* and was included for comparison. Phylogenetic analysis supports that these genes form a paralogous pair in *Z. palustris* (Figure 4F). Phylogenetic analysis of *SRG1* homologs in monocot grasses revealed that *ZPchr0001g31674* clustered with its *Z. palustris* paralog*, ZPchr0001g29384*, followed by *Zizania latifolia* within the Oryzeae clade. *Zizania* homologs were most similar to *Oryza brachyantha* and *Oryza meyeriana*. Protein sequence alignment between the two *Zizania* paralogs showed two amino acid substitutions, and a 34-residue insertion present only in *ZPchr0001g31674*. Domain annotation indicated that this insertion overlaps with the C-terminal region of the 2OG-Fe(II) oxygenase domain, while in *ZPchr0001g29384* the domain terminates upstream of the insertion site (Figure S3). Expression data (Figure 4e) show that transcript levels of *ZPchr0001g29384* are higher than *ZPchr0001g31674* across all tissues, and it is differentially expressed only in the flag leaf versus aerial leaf comparison.

Across all three transitions, leaf development in *Z. palustris* progresses from large-scale transcriptomic reprogramming during aquatic-to-aerial emergence, to selective remodeling of hormone and wall-associated pathways in early aerial growth, and finally to suppression of structural programs with activation of metabolic processes in the flag leaf. This progression delineates the molecular sequence by which leaves transition from juvenile aquatic forms to structurally mature, metabolically active organs at the reproductive stage.

### 3.5 Floral Transcriptome Differentiation and Expression of Seed Shattering Genes

To better understand sex-specific floral development in *Z. palustris*, we conducted transcriptome profiling of unisexual panicles representing staminate (male) and pistillate (female) organs. This developmental differentiation is central to reproductive success and grain production in this outcrossing species, where pollen and ovule development occur on separate flowers.

Between male and female florets, 1,078 genes were differentially expressed, with 692 upregulated in male panicles and 386 in female panicles (Figure S4). Upregulated genes in female and male panicles, relative to each other, were enriched for 11 and 66 GO terms, respectively (Table S8). Female panicles showed increased expression of genes associated with mitotic progression and hormone homeostasis, including *Cyclin A2;1* (*ZPchr0009g121*), Cytokinin oxidase/dehydrogenase (*ZPchr0008g12028*), and Growth-regulating factor 9 (*ZPchr0003g17492*) (Figure S4).

Male panicles were enriched for transcriptional programs associated with reproductive development, including MADS-box transcription factor 16 (*ZPchr0010g10752*), Pollen receptor-like kinase 3 (*ZPchr0011g27156*), and Phl p 11-like allergen (*ZPchr0008g14104*) (Figure S4). Additional male-biased expression was detected for genes involved in amino acid metabolism, biogenic amine biosynthesis, and energy reserve mobilization (Table S8). Auxin-associated genes were also upregulated, including *YUCCA10 (ZPchr0011g28119), ARF2 (ZPchr0001g32063*), and enzymes involved in indole biosynthesis *(ZPchr0002g23314, ZPchr0008g12535, ZPchr0013g35689)* (Figure S4). These were enriched in indolalkylamine biosynthetic processes, suggesting auxin metabolism plays a key role in male floral development. Genes associated with cuticle biosynthesis *(ZPchr0010g8337, ZPchr0010g9722, ZPchr0009g200)* and pathogen defense *(ZPchr0005g14516, ZPchr0005g15165, ZPchr0010g9678)* were also more highly expressed in male panicles, consistent with enhanced structural protection and stress responses in staminate tissues.

Given the importance of floral architecture and grain retention for domestication, we examined the expression of known *O. sativa* shattering regulators as seed shattering remains a major limitation in *Z. palustris* grain production, contributing to substantial yield loss. Comparative genomics identified *Z. palustris* homologs of *qSH1, SH1, SHAT1, SH4, SH5,* and *LG1* (Haas et al., 2021; Millas et al., 2025). Many of these genes occurred as duplicate or triplicate copies, consistent with the whole-genome duplication and subsequent repetitive element expansion in *Z. palustris* (Haas et al., 2021). Additional homologs were identified in this study, including *SLR1, NPC1, RHS1, SNB, GRF4,* and *SH11* (Table 1).

**Table 1.**
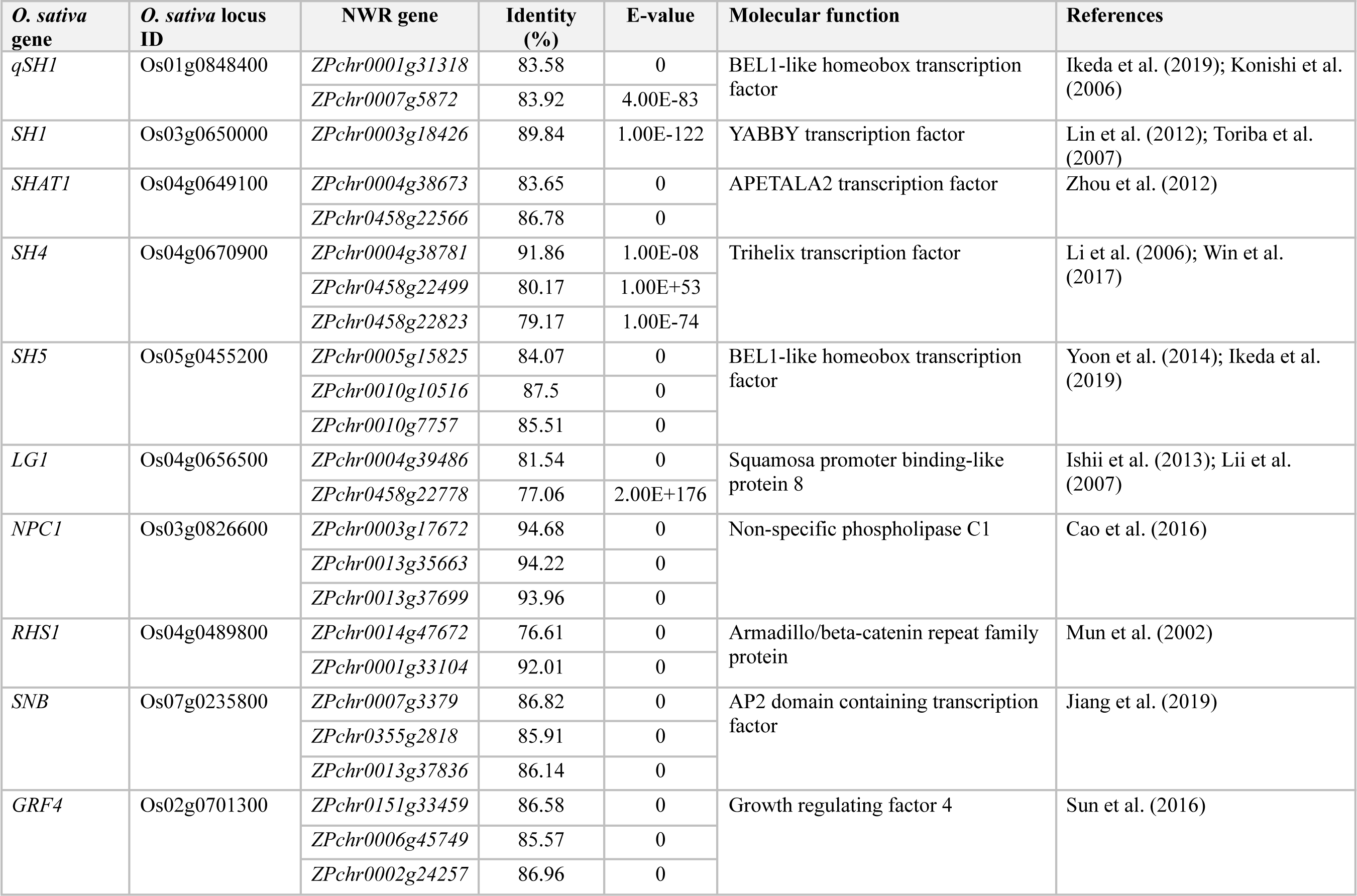
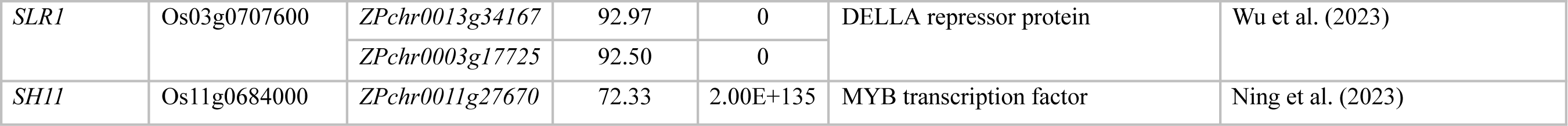
List of *Z. palustris* genes orthologous to *O. sativa* seed shattering related genes.

Expression profiling revealed that many shattering-related homologs displayed tissue-specific or paralog-specific expression, with most showing peak expression in vegetative tissues or early reproductive structures rather than in panicles (Figure 5). For instance, the two *qSH1* paralogs, *ZPchr0001g31318* and *ZPchr0007g5872*, were most highly expressed in roots and stem first nodes, respectively. Among the three *SH5* paralogs, expressions diverged widely, peaking in flag leaves (*ZPchr0010g7757*), stem first nodes (*ZPchr0010g10516*), and roots (*ZPchr0005g15825*). Only a subset of shattering genes showed elevated expression in reproductive tissues, including one *LG1* paralog (*ZPchr0458g22778*) in female panicles and one *SNB* paralog (*ZPchr0007g3379*) in male panicles. By contrast, multiple shattering homologs (*SH1, qSH1, SH5, SH4, SHAT1, NPC1, RHS1, SNB, GRF4,* and *SH11*) displayed strong expression in stems during early and late booting stages. These findings suggest subfunctionalization or regulatory divergence among paralogs, highlighting a subset of candidates for future functional analysis of seed shattering in *Z. palustris*.

**Figure 5.**
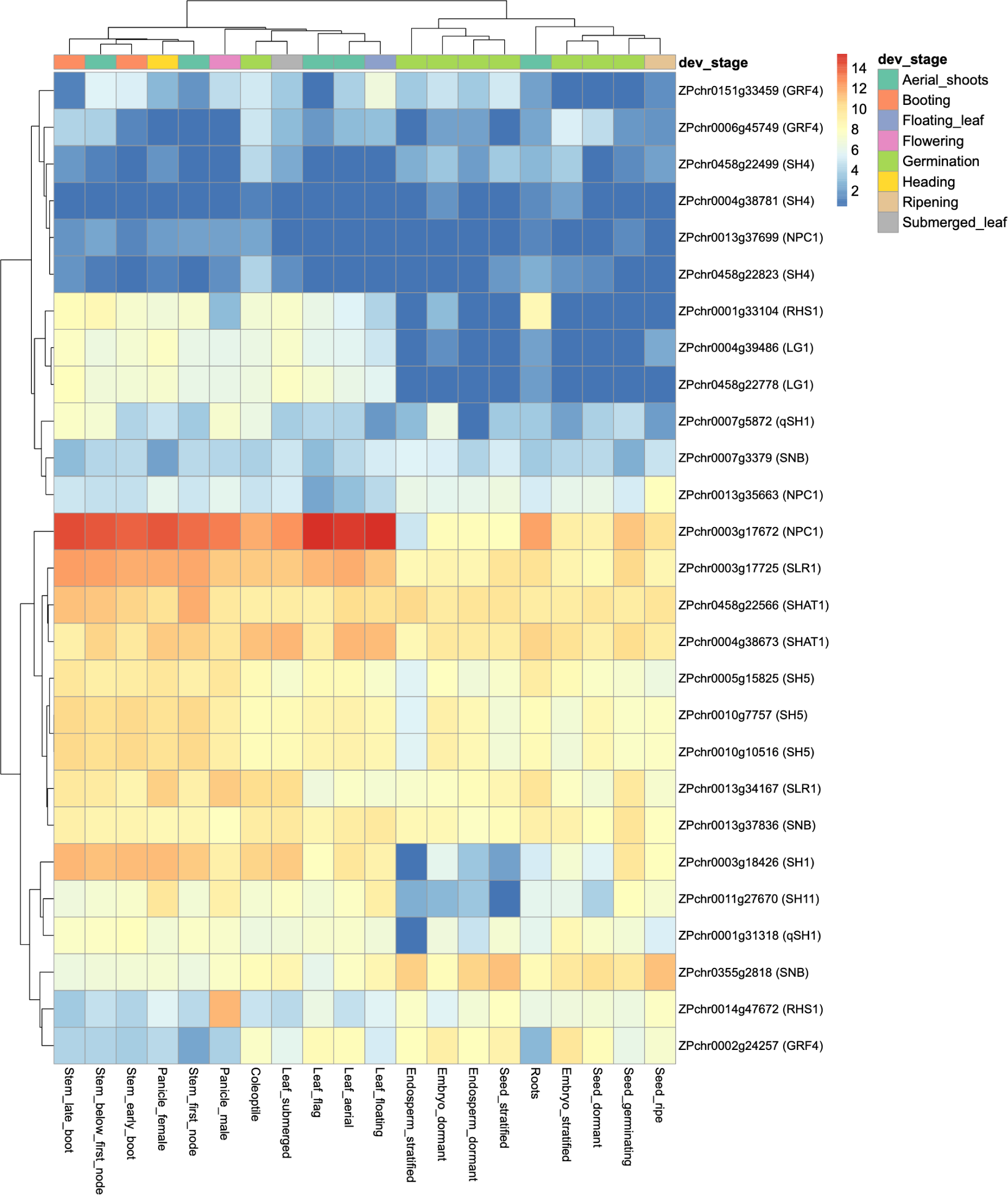
Expression of seed shattering orthologs across *Zizania palustris* tissues. Heatmap of variance-stabilized expression (vst) for orthologs of known *Oryza sativa* shattering genes across 20 tissues. Tissues span vegetative, reproductive, and seed developmental stages.

## 4 Discussion

This study presents the first transcriptomic atlas spanning the full annual life cycle of *Z. palustris*, a grain-producing, aquatic grass native to North America that is both ecologically embedded in the shallow freshwater of the Great Lakes region and cultivated commercially in man-made, irrigated paddies. As a crop wild relative of rice that diverged 20–30 MYA and experienced lineage-specific genome duplication, *Z. palustris* provides a powerful complement to *Oryza* as a comparative system while capturing ecological dimensions largely absent from terrestrial cereals, including seasonal submergence, anaerobic germination, aquatic juvenile growth, and obligate outcrossing with unisexual inflorescences. Our results highlight both validating findings consistent with previous studies in *Oryza* and Arabidopsis as well as novel insights unique to *Zizania*. Together with the recent reference genome (Haas et al., 2021), this atlas links gene models to their tissue and stage expression, enabling candidate-gene discovery and functional annotation for traits relevant to domestication, adaptation, and conservation in *Zizania*.

Seed development in *Z. palustris* revealed transcriptional changes in ABA- and GA-associated pathways that align with its strong physiological dormancy. Elevated expression of ABA biosynthetic genes such as *NCED* and *CYP707A* and the concurrent repression of GA biosynthetic enzymes (*GA20ox*) suggest a transcriptional state associated with dormancy maintenance, consistent with classical models in cereals (Seo et al., 2006; Nonogaki, 2019). Shifts in DELLA proteins and ABA signaling components suggest coordinated hormone-related transcriptional programs that coincide with endosperm maturation and embryo quiescence, which have been linked to stratification-dependent germination, as described in rice and barley (Rodríguez-Gacio et al., 2009; Nonogaki, 2019). In *Z. palustris*, persistence of this transcriptional program likely reflects adaptation for prolonged cold exposure before germination, a trait that promotes staggered emergence in wetlands but hinders uniform establishment in cultivation. These transcriptomic patterns are consistent with prior physiological observations that chemical scarification or exogenous hormones fail to release dormancy in freshly harvested *Z. palustris* seeds (McGilp et al., 2022). Beyond hormonal regulation, mitochondrial and respiratory genes peaked during mid-seed development, marking the transition from anabolic storage programs to catabolic energy management, a feature that may be particularly important for seeds maturation in aquatic, low-oxygen environments (Finch-Savage and Bassel, 2016). These findings highlight the endosperm as a key transcriptional hub integrating hormone-related and metabolic reprogramming during dormancy and germination transitions, reflecting its ecological adaptation to wetland habitats and a critical consideration for domestication of *Z. palustris*.

Leaf development highlighted a stress-related progression from hypoxic submerged juvenile tissues to photosynthetically competent flag leaves. Submerged leaves strongly induced hypoxia-responsive and anaerobic metabolism genes, consistent with adaptations to early vegetative growth underwater (Mustroph et al., 2010). Floating leaves exhibited transient activation of oxidative stress regulators, including expansins (EXPA), TCP-domain transcription factors, and photosynthetic components, indicating coordinated cell wall remodeling and energy capture. The regulation of *GIL1* and vesicle-trafficking genes such as *RET3* suggests a coordinated mechanism for cuticle deposition and leaf structural integrity as leaves transition to aerial growth. In flag leaves, transcriptional profiles are dominated by photosynthesis and carbohydrate-metabolism pathways. These transitions reflect a progressive rebalancing of cellular redox homeostasis, with ROS-scavenging enzymes and hormone pathways (ethylene, jasmonate) mitigating the costs of reoxygenation, a process well documented in rice and other wetland plants where reoxygenation stress is a critical bottleneck (Fukao & Bailey-Serres, 2008). Here, we show at transcriptome scale how an emergent aquatic cereal resolves this challenge. For crop improvement, optimizing this transition offers avenues to enhance resilience to flooding or fluctuating water tables; for wetland management, it highlights the fine-scale mechanisms that enable persistence under dynamic hydrological conditions.

Reproductive development revealed both conserved and novel regulators of shattering in *Z. palustris*. We identified high confidence orthologs of rice shattering genes and found widespread evidence of paralog subfunctionalization, likely driven by ancestral duplication and genome expansion events (Haas et al., 2021). This redundancy likely contributes to the incomplete suppression of shattering in domesticated lines, which complicates mechanical harvest. Recent studies in rice have also positioned shattering as a polygenic, stepwise process, in which coordinated modulation of multiple QTLs, panicle architecture, hormone signaling, and lignin biosynthesis collectively determine seed retention (Ishikawa et al., 2022; Maity et al., 2021). Our transcriptome data suggests that *Z. palustris* may follow a similarly complex architecture, where multiple paralogs of *qSH1, SH4, SH5*, and other shattering-related genes exhibit spatiotemporally distinct expression, particularly in stems, booting tissues, and reproductive organs. This layered expression pattern is consistent with a quantitative model in which partial expression reductions across several loci collectively reduce abscission strength, rather than relying on a single gene knockout. The persistence of multiple functional copies in *Z. palustris* suggests that more precise interventions, whether allele selection, targeted editing, or regulatory tuning, will be needed to achieve durable seed retention. This case illustrates how domestication in orphan crops may diverge from canonical models in cereals and underscores the value of transcriptomic resolution for prioritizing gene targets.

Viewed broadly, *Z. palustris* illustrates the value of underutilized crops as climate-resilient and ecologically integrated food sources, and as tractable comparative systems. Its life cycle that naturally interrogates hypoxia–reoxygenation biology, its strong synteny with *Oryza* enables efficient candidate transfer, and its immediate translational value for wetland stewardship and culturally important foods. The atlas defines stable housekeeping sets, tissue-resolved gene modules, and high-priority candidates for domestication traits such as shattering and maturation synchrony, providing a platform for GWAS, eQTL mapping, and other validation studies in a crop wild relative poised for responsible improvement. By situating *Z. palustris* at the intersection of ecological adaptation and domestication, this work underscores the potential of transcriptomics to accelerate the improvement of orphan crops while deepening our understanding of plant diversification in dynamic environments.

## Abbreviations

ABA: abscisic acid
CK: cytokinin
CoV: coefficient of variation
cNWR: cultivated Northern Wild Rice
CWR: crop wild relative
DEGs: differentially expressed genes
eQTL: expression quantitative locus
GA: gibberellin
GO: Gene Ontology
GWAS: genome-wide association study
HK: housekeeping genes
JA: jasmonic acid
MFC: minimum fold change
MYA: million years ago
NWR: Northern Wild Rice
TPM: transcripts per million
WGD: whole genome duplication.

## Acknowledgments

The authors acknowledge financial support from the State of Minnesota Agricultural Research, Education, Extension, and Technology Transfer Program. The authors also thank the University of Minnesota Genomics Center for sequencing support and the Minnesota Supercomputing Institute for providing computational resources that enabled data analysis and storage.

## DATA AVAILABILITY

Raw sequencing reads have been deposited in the NCBI BioProject under the accession number PRJNA1301937. The functional annotations, TPM expression matrices at both the transcript and gene levels are publicly available from the Figshare repository (https://doi.org/10.6084/m9.figshare.30018808).

## Supplemental Material

The supplemental material includes tables summarizing gene expression levels across 20 tissues, hormone-related differential expression during seed dormancy transitions, and gene ontology enrichment analyses across developmental stages in Zizania palustris. Additional datasets provide RNA-seq data used to construct the gene expression atlas, cluster-level functional enrichment, tissue-specificity metrics, and genes with stable expression profiles, along with figures illustrating the life cycle of Northern Wild Rice, tissue-specific expression patterns, protein domain annotations, and sex-specific expression in panicles.

## References

Alexa, A., & Rahnenfuhrer, J. (2016). topGO: Enrichment analysis for gene ontology. R package version 2.46.0. Available at https://bioconductor.org/packages/topGO

Allen, T., Ingles, P. J., Praekelt, U., Smith, H., & Whitelam, G. C. (2006). Phytochrome-mediated agravitropism in Arabidopsis hypocotyls requires *GIL1* and confers a fitness advantage. The Plant Journal, 46(4), 641–648. 10.1111/j.1365-313X.2006.02727.x

Andrews, S. (2010). FastQC: a quality control tool for high throughput sequence data. Available at http://www.bioinformatics.babraham.ac.uk/projects/fastqc

Bolger, A. M., Lohse, M., & Usadel, B. (2014). Trimmomatic: A flexible trimmer for Illumina sequence data. Bioinformatics, 30(15), 2114–2120. 10.1093/bioinformatics/btu170

Bushnell, B. (2014). BBTools. Available at https://sourceforge.net/projects/bbmap/

Cho, H.-T., & Kende, H. (1997). Expression of Expansin Genes Is Correlated with Growth in Deepwater Rice. The Plant Cell, 9(9), 1661–1671. 10.2307/3870451

Danecek, P., Bonfield, J. K., Liddle, J., Marshall, J., Ohan, V., Pollard, M. O., Whitwham, A., Keane, T., McCarthy, S. A., Davies, R. M., & Li, H. (2021). Twelve years of SAMtools and BCFtools. GigaScience, 10(2), giab008. 10.1093/gigascience/giab008

Duquette, J., & Kimball, J. A. (2020). Phenological stages of cultivated northern wild rice according to the BBCH scale. Annals of Applied Biology, 176(3), 350–356. 10.1111/aab.12588

Eddy, S. R. (2011). Accelerated Profile HMM Searches. PLoS Computational Biology, 7(10), e1002195. 10.1371/journal.pcbi.1002195

Finch-Savage, W. E., & Bassel, G. W. (2016). Seed vigour and crop establishment: extending performance beyond adaptation. Journal of Experimental Botany, 67(3), 567–591.

Fukao, T., & Bailey-Serres, J. (2008). Ethylene—a key regulator of submergence responses in rice. Plant Science, 175(1–2), 43–51.

Guo, L., Qiu, J., Han, Z., Ye, Z., Chen, C., Liu, C., Xin, X., Ye, C., Wang, Y., Xie, H., Wang, Y., Bao, J., Tang, S., Xu, J., Gui, Y., Fu, F., Wang, W., Zhang, X., Zhu, Q., … Fan, L. (2015). A host plant genome (*Zizania latifolia*) after a century-long endophyte infection. The Plant Journal, 83(4), 600–609. 10.1111/tpj.12912

Guo, Y., & Ge, S. (2005). Molecular phylogeny of Oryzeae (Poaceae) based on DNA sequences from chloroplast, mitochondrial, and nuclear genomes. American Journal of Botany, 92(9), 1548–1558. 10.3732/ajb.92.9.1548

Haas, B. J. (2019). TransDecoder (Find Coding Regions Within Transcripts). Version 5.7.1. Available at https://github.com/TransDecoder/TransDecoder/wiki

Haas, M., Kono, T., Macchietto, M., Millas, R., McGilp, L., Shao, M., Duquette, J., Qiu, Y., Hirsch, C. N., & Kimball, J. (2021). Whole-genome assembly and annotation of northern wild rice, *Zizania palustris* L., supports a whole-genome duplication in the *Zizania* genus. The Plant Journal, 107(6), 1802–1818. 10.1111/tpj.15419

Ishikawa, R., Castillo, C. C., Htun, T. M., Numaguchi, K., Inoue, K., Oka, Y., … & Ishii, T. (2022). A stepwise route to domesticate rice by controlling seed shattering and panicle shape. Proceedings of the National Academy of Sciences, 119(26), e2121692119.

Jin, H. (2000). Transcriptional repression by AtMYB4 controls production of UV-protecting sunscreens in Arabidopsis. The EMBO Journal, 19(22), 6150–6161. 10.1093/emboj/19.22.6150

Jones, P., Binns, D., Chang, H.-Y., Fraser, M., Li, W., McAnulla, C., McWilliam, H., Maslen, J., Mitchell, A., Nuka, G., Pesseat, S., Quinn, A. F., Sangrador-Vegas, A., Scheremetjew, M., Yong, S.-Y., Lopez, R., & Hunter, S. (2014). InterProScan 5: Genome-scale protein function classification. Bioinformatics, 30(9), 1236–1240. 10.1093/bioinformatics/btu031

Kassambara, A., & Mundt, F. (2020). factoextra: Extract and visualize the results of multivariate data analyses. R package version 1.0.7. Available at https://CRAN.R-project.org/package=factoextra

Kim, D., Langmead, B., & Salzberg, S. L. (2015). HISAT: A fast spliced aligner with low memory requirements. Nature Methods, 12(4), 357–360. 10.1038/nmeth.3317

Kolde, R. (2019). pheatmap: Pretty Heatmaps. R package version 1.0.12. Available at https://CRAN.R-project.org/package=pheatmap

Letunic, I., Khedkar, S., & Bork, P. (2021). SMART: Recent updates, new developments and status in 2020. Nucleic Acids Research, 49(D1), D458–D460. 10.1093/nar/gkaa937

Li, G., Zhang, J., Li, J., Yang, Z., Huang, H., & Xu, L. (2012). Imitation Switch chromatin remodeling factors and their interacting RINGLET proteins act together in controlling the plant vegetative phase in Arabidopsis. The Plant Journal, 72(2), 261–270. 10.1111/j.1365-313X.2012.05074.x

Love, M. I., Huber, W., & Anders, S. (2014). Moderated estimation of fold change and dispersion for RNA-seq data with DESeq2. Genome Biology, 15(12), 550. 10.1186/s13059-014-0550-8

Lüleci, H. B., & Yılmaz, A. (2022). Robust and rigorous identification of tissue-specific genes by statistically extending tau score. BioData Mining, 15(1), 31. 10.1186/s13040-022-00315-9

Machado, F. B., Moharana, K. C., Almeida-Silva, F., Gazara, R. K., Pedrosa-Silva, F., Coelho, F. S., Grativol, C., & Venancio, T. M. (2020). Systematic analysis of 1298 RNA-Seq samples and construction of a comprehensive soybean (Glycine max) expression atlas. The Plant Journal, 103(5), 1894–1909. 10.1111/tpj.14850

Maity, A., Lamichaney, A., Joshi, D. C., Bajwa, A., Subramanian, N., Walsh, M., & Bagavathiannan, M. (2021). Seed shattering: a trait of evolutionary importance in plants. Frontiers in Plant Science, 12, 657773.

McGilp, L., Castell-Miller, C., Haas, M., Millas, R., & Kimball, J. (2023). Northern Wild Rice (*Zizania palustris* L.) breeding, genetics, and conservation. Crop Science, 63(4), 1904–1933. 10.1002/csc2.20973

Mi, H., Ebert, D., Muruganujan, A., Mills, C., Albou, L.-P., Mushayamaha, T., & Thomas, P. D. (2021). PANTHER version 16: A revised family classification, tree-based classification tool, enhancer regions and extensive API. Nucleic Acids Research, 49(D1), D394–D403. 10.1093/nar/gkaa1106

Millas, R., McGilp, L., Mickelson, A., Banting, M., Castell-Miller, C., & Kimball, J. (2025). Seed Shattering in a North American Oryzeae grain: Developmental and Genomic Signatures of Early Domestication. 10.21203/rs.3.rs-7032638/v1

Mistry, J., Chuguransky, S., Williams, L., Qureshi, M., Salazar, G. A., Sonnhammer, E. L. L., Tosatto, S. C. E., Paladin, L., Raj, S., Richardson, L. J., Finn, R. D., & Bateman, A. (2021). Pfam: The protein families database in 2021. Nucleic Acids Research, 49(D1), D412–D419. 10.1093/nar/gkaa913

Mitchum, M. G., Yamaguchi, S., Hanada, A., Kuwahara, A., Yoshioka, Y., Kato, T., Tabata, S., Kamiya, Y., & Sun, T. (2006). Distinct and overlapping roles of two gibberellin 3-oxidases in Arabidopsis development. The Plant Journal, 45(5), 804–818. 10.1111/j.1365-313X.2005.02642.x

Mustroph, A., Lee, S. C., Oosumi, T., Zanetti, M. E., Yang, H., Ma, K., … & Bailey-Serres, J. (2010). Cross-kingdom comparison of transcriptomic adjustments to low-oxygen stress highlights conserved and plant-specific responses. Plant Physiology, 152(3), 1484–1500.

Mustroph, A., Sonnewald, U., & Biemelt, S. (2007). Characterisation of the ATP-dependent phosphofructokinase gene family from *Arabidopsis thaliana*. FEBS Letters, 581(13), 2401–2410. 10.1016/j.febslet.2007.04.060

Nonogaki, H. (2019). Seed germination and dormancy: The classic story, new puzzles, and evolution. Journal of Integrative Plant Biology, 61(5), 541–563.

Pence, V. C. (1995). Cryopreservation of recalcitrant seeds. In Y. P. S. Bajaj (Ed.), Biotechnology in Agriculture and Forestry 32, Cryopreservation of Plant Germplasm (Vol. 1, pp. 29–50).

Pérez-Pérez, J. M., Esteve-Bruna, D., González-Bayón, R., Kangasjärvi, S., Caldana, C., Hannah, M. A., Willmitzer, L., Ponce, M. R., & Micol, J. L. (2013). Functional Redundancy and Divergence within the Arabidopsis RETICULATA-RELATED Gene Family. Plant Physiology, 162(2), 589–603. 10.1104/pp.113.217323

Pertea, M., Pertea, G. M., Antonescu, C. M., Chang, T.-C., Mendell, J. T., & Salzberg, S. L. (2015). StringTie enables improved reconstruction of a transcriptome from RNA-seq reads. Nature Biotechnology, 33(3), 290–295. 10.1038/nbt.3122

Phan, H. A., Iacuone, S., Li, S. F., & Parish, R. W. (2011). The MYB80 Transcription Factor Is Required for Pollen Development and the Regulation of Tapetal Programmed Cell Death in *Arabidopsis thaliana*. The Plant Cell, 23(6), 2209–2224. 10.1105/tpc.110.082651

Pruneda-Paz, J. L., Breton, G., Para, A., & Kay, S. A. (2009). A Functional Genomics Approach Reveals CHE as a Component of the *Arabidopsis* Circadian Clock. Science, 323(5920), 1481–1485. 10.1126/science.1167206

Quast, C., Pruesse, E., Yilmaz, P., Gerken, J., Schweer, T., Yarza, P., Peplies, J., & Glöckner, F. O. (2012). The SILVA ribosomal RNA gene database project: Improved data processing and web-based tools. Nucleic Acids Research, 41(D1), D590–D596. 10.1093/nar/gks1219

Rodríguez-Gacio, M. D. C., Matilla-Vázquez, M. A., & Matilla, A. J. (2009). Seed dormancy and ABA signaling: the breakthrough goes on. Plant Signaling & Behavior, 4(11), 1035–1048.

Seo, M., Hanada, A., Kuwahara, A., Endo, A., Okamoto, M., Yamauchi, Y., … & Nambara, E. (2006). Regulation of hormone metabolism in Arabidopsis seeds: phytochrome regulation of abscisic acid metabolism and abscisic acid regulation of gibberellin metabolism. The Plant Journal, 48(3), 354–366.

Signal, B., & Kahlke, T. (2022). how_are_we_stranded_here: Quick determination of RNA-Seq strandedness. BMC Bioinformatics, 23(1), 49. 10.1186/s12859-022-04572-7

Soneson, C., Love, M. I., & Robinson, M. D. (2015). Differential analyses for RNA-seq: Transcript-level estimates improve gene-level inferences. F1000Research, 4, 1521. 10.12688/f1000research.7563.1

Staswick, P. E., Serban, B., Rowe, M., Tiryaki, I., Maldonado, M. T., Maldonado, M. C., & Suza, W. (2005). Characterization of an Arabidopsis Enzyme Family That Conjugates Amino Acids to Indole-3-Acetic Acid. The Plant Cell, 17(2), 616–627. 10.1105/tpc.104.026690

Tanaka, K., Murata, K., Yamazaki, M., Onosato, K., Miyao, A., & Hirochika, H. (2003). Three Distinct Rice Cellulose Synthase Catalytic Subunit Genes Required for Cellulose Synthesis in the Secondary Wall. Plant Physiology, 133(1), 73–83. 10.1104/pp.103.022442

Wickham, H. (2016). ggplot2: Elegant Graphics for Data Analysis. Springer-Verlag, New York. ISBN 978-3-319-24277-4. Available at https://ggplot2.tidyverse.org

Yan, N., Yang, T., Yu, X.-T., Shang, L.-G., Guo, D.-P., Zhang, Y., Meng, L., Qi, Q.-Q., Li, Y.-L., Du, Y.-M., Liu, X.-M., Yuan, X.-L., Qin, P., Qiu, J., Qian, Q., & Zhang, Z.-F. (2022). Chromosome-level genome assembly of *Zizania latifolia* provides insights into its seed shattering and phytocassane biosynthesis. Communications Biology, 5(1), 36. 10.1038/s42003-021-02993-3

Zhong, R., Lee, C., Zhou, J., McCarthy, R. L., & Ye, Z.-H. (2008). A Battery of Transcription Factors Involved in the Regulation of Secondary Cell Wall Biosynthesis in *Arabidopsis*. The Plant Cell, 20(10), 2763–2782. 10.1105/tpc.108.061325

